# Trait dimensionality in experimental and natural ecological communities

**DOI:** 10.64898/2026.03.10.710797

**Authors:** Vicente J. Ontiveros, David Alonso, José A. Capitán

**Affiliations:** Complex Systems Group, Department of Applied Mathematics, Universidad Politécnica de Madrid, Spain; Ecology and Complexity. Center for Advanced Studies of Blanes (CEAB-CSIC), Blanes, Catalonia, Spain

## Abstract

Classical niche theory predicts that species with more similar traits experience stronger competition, and thus, trait dimensionality, here defined as the number of independent traits per species in a community, plays a critical role in coexistence. Despite this, current studies often assume that communities are structured by only a few key traits. Here, we leveraged a theoretical framework that integrates phylogenetic information with repeated instances of community assembly to estimate the number of ecologically relevant traits required to support observed species richness in experimental and natural plant communities. We found that the inferred trait dimensionality is surprisingly high, often exceeding species richness, suggesting that many traits contribute to coexistence. Furthermore, we explored drivers of grassland trait dimensionality, and it depends in complex ways on area, species pool size, and latitude. Our findings indicate that local coexistence may rely on a larger number of traits than previously assumed, challenging low-dimensional trait-based views of community structure.

## 1 Introduction

Competition has always played a prominent role in our understanding of the structure of ecological communities. In *On the Origin of Species*, Darwin noted that “competition will generally be most severe between those forms which are most nearly related to each other in habits, constitution, and structure.” Building on this idea, the niche concept –defined as a multidimensional hypervolume [1] that encapsulates a species’ ecological and functional characteristics– provides a framework for understanding species coexistence. To study the niche, ecologists typically measure traits and categorize them into distinct dimensions that align with human reasoning [2]. However, the fundamental dimensionality of these trait spaces has not been explored through theoretical predictions that explicitly incorporate evolutionary history (phylogenies), coexistence, and competition. Here, we estimate the trait dimensionality that allows for a certain species richness in ecological communities given their species pool and shared evolutionary history. Using both experimental and observational data, we show that, with some corrections, mean field competition can be a good predictor of coexistence, and trait dimensionality might be higher than previously recognized.

Previous studies have attempted to quantify the dimensionality of ecological systems. In a pioneering study, less than 10 dimensions were necessary to fully describe ecological networks, including food-webs, antagonistic, and mutualistic networks [3]. Another foundational study indicates that the plant trait space is restricted to about 6 dimensions [4]. These initial studies catalyzed further analyses, which found low dimensionality, in plants [5, 6, 7] and other organisms, such as birds [8, 9], ants [10], fishes [2], or mammals [9]. Moreover, a recent synthesis over a wide range of organisms identifies a typical dimensionality of the trait space of 3-6 dimensions [11]. However, several studies point out the limitations of trait dimensionality to explain local community assembly. Although conceptualizing trait spaces in comprehensible dimensions have important advantages, at local scales other factors influence trait combinations [6, 9], and may even render the trait dimension paradigm unapplicable [12]. Yet, this low dimensionality raises a fundamental question about species coexistence. If the trait space is so constrained, how can that many species live in the same place [13]?

Phylogeny plays a fundamental role in shaping community assembly, influencing patterns of species coexistence and competitive interactions [14]. Research in this area has provided deep insights into the mechanisms underlying this influence. For instance, [15] demonstrated that competitive exclusion can lead to phylogenetic overdispersion when differences in competitive ability are small, whereas strong competitive differences correlated with phylogenetic distance can instead drive phylogenetic clustering. Phylogenetic structure also informs predictions of species coexistence and abundance patterns, as shown recently through consumer-resource models [16]. Further theoretical developments extend these insights by integrating trait evolution with Lotka-Volterra dynamics, offering a quantitative framework to assess community properties and estimate trait dimensionality [17].

We build on theoretical expectations [17] to estimate trait dimensionality under the assumption of equal competitive abilities. In this context, we treat traits as idealized functional properties, representing measurable aspects of an organism, such as morphology, physiology, or behavior. While abstract, these traits are estimated in specific ecological contexts, which aligns them conceptually with niche dimensions. This interpretation allows trait-based analyses to serve as a proxy for niche structure, especially when direct measurements are unavailable. Here, we assess trait dimensionality in both experimental (grassland) and natural (grassland and forest) communities, testing whether similar communities exhibit comparable trait distribution patterns. As an additional consistency check, we explore the key factors influencing dimensionality. Furthermore, we evaluate whether a simplified phylogeny, representing mean-field competition, can effectively capture trait dimensionality. Our findings indicate that dimensionality is higher than previously recognized.

## 2 Material & Methods

### 2.1 Theoretical framework

The estimation of the effective number of traits, *ℓ*, that match an expected diversity if based on the theoretical analyses presented in reference [17]. Here, we provide a summary of the procedure used to carry out that estimation. The rationale behind this procedure is summarized in figure 1.

**Figure 1.**
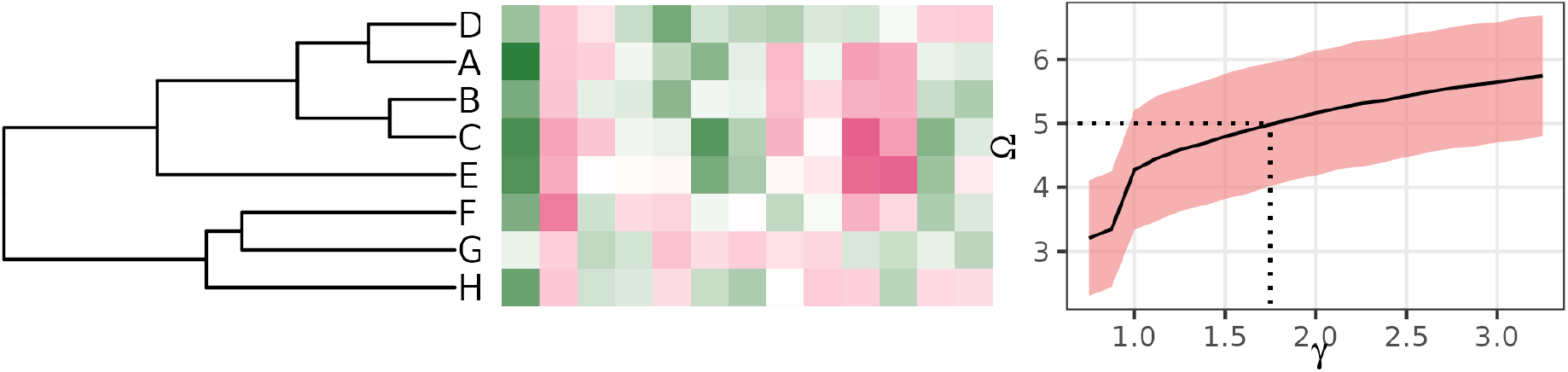
Framework to estimate trait dimensionality. *Left*, from an empirical phylogeny, we extract the shared evolutionary history between species. *Middle*, a variable number of traits reflect a phylogenetic random walk based on the shared history, and consequently determine an expected species richness. *Right*, multiple realizations of the random walk allow for the estimation of trait dimensionality given the observed species richness of the community.

We start with an empirical phylogeny, from which we derive the associated variance-covariance matrix, Σ, where each element Σ_*ij*_ represents the shared evolutionary history (branch length) between species *i* and *j*. Diagonal elements are Σ_*ii*_ = 1 for all *i*. Off-diagonal elements are obtained as Σ_*ij*_ = 1 *− t*_*ij*_, where *t*_*ij*_ is the time (in units in which the total length of the tree is one) at which species *i* adn *j* coalesce to their common ancestor.

This evolutionary history shapes interactions between species, mediated through a specified set of traits, *ℓ*. Assuming evolution through diffusion processes [18], these traits follow a multivariate normal distribution with the covariance matrix Σ generated by the tree. So, we sample the specific multivariate normal distribution to obtain a species-by-trait matrix *G*, whose rows are the vectors of traits sampled for each species. Then, assuming stronger competition for species that have similar trait vectors, we construct an interaction matrix as 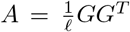, measuring species interactions proportionally to trait vector overlap. Given that *Q* is formed by samples of a multivariate normal, matrix *A* (of size *n × n*) is therefore a sample of the Wishart ensemble, *A* ~ *W*_*n*_(*ℓ*^−1^Σ, *ℓ*) with *ℓ* degrees of freedom. Therefore, given the empirical phylogeny (parameterized by the tree covariance matrix Σ), we draw interaction matrices by sampling from the Wishart distribution.

Using this interaction matrix within a Generalized Lotka-Volterra (GLV) model, one can show that the dynamics tends asymptotically to a globally stable attractor, in which some of the species survive and some others do not [19]. The size of the attractor can be determined by solving a linear programming problem (see [17] and the online repository associated to that reference, [20]). Determining the size fo the attractor (i.e., the number of survivors predicted by GLV model, given the empirical tree used to construct species interactions) can be done through the Lemke-Howson algorithm. The reader is referred to reference [17] for further details.

We identify the system’s attractor, which determines the equilibrium number of surviving species. Different samples of the interaction matrix yield varying sizes of the attractors, for fixed *ℓ*: averaing attractor’s size over the noise induced by the interactions yield the confidence intervals shown in figure 1 (right panel). Importantly, the number of survivors depends on the dimensionality of trait space, *ℓ*, which influences the generated interactions. By adjusting *ℓ* to match an observed diversity Ω, we can identify the value that yields a number of survivor species closest to the observed number derived from the phylogenetic tree (figure 1, right panel).

### 2.2 Datasets

To evaluate whether the currently measured trait dimensionality is sufficient to explain species coexistence, we analyzed two datasets containing information on species traits, replication of community assembly, and a definite species list. The first dataset [21] corresponds to a outdoor mesocosm experiment of community assembly of 34 floodplain plant species over three years. The authors measured or extracted from databases 45 plant traits for these

Temperate grasslands are the predominant habitat with experimental data on community assembly and readily available phylogenies. Such data is surprisingly scarce and comes mainly from BEF experiments, but may be extracted for several other sources as mesocosms, common garden, or community assembly experiments. Table 1 summarises the grassland experiments used in this manuscript. We selected studies that had a well-defined species pool with at least eight species, included plot replication, provided data on final species richness, and featured control or minimally manipulated plots. All experiments were conducted at latitudes between 37° and 57°N.

**Table 1:**
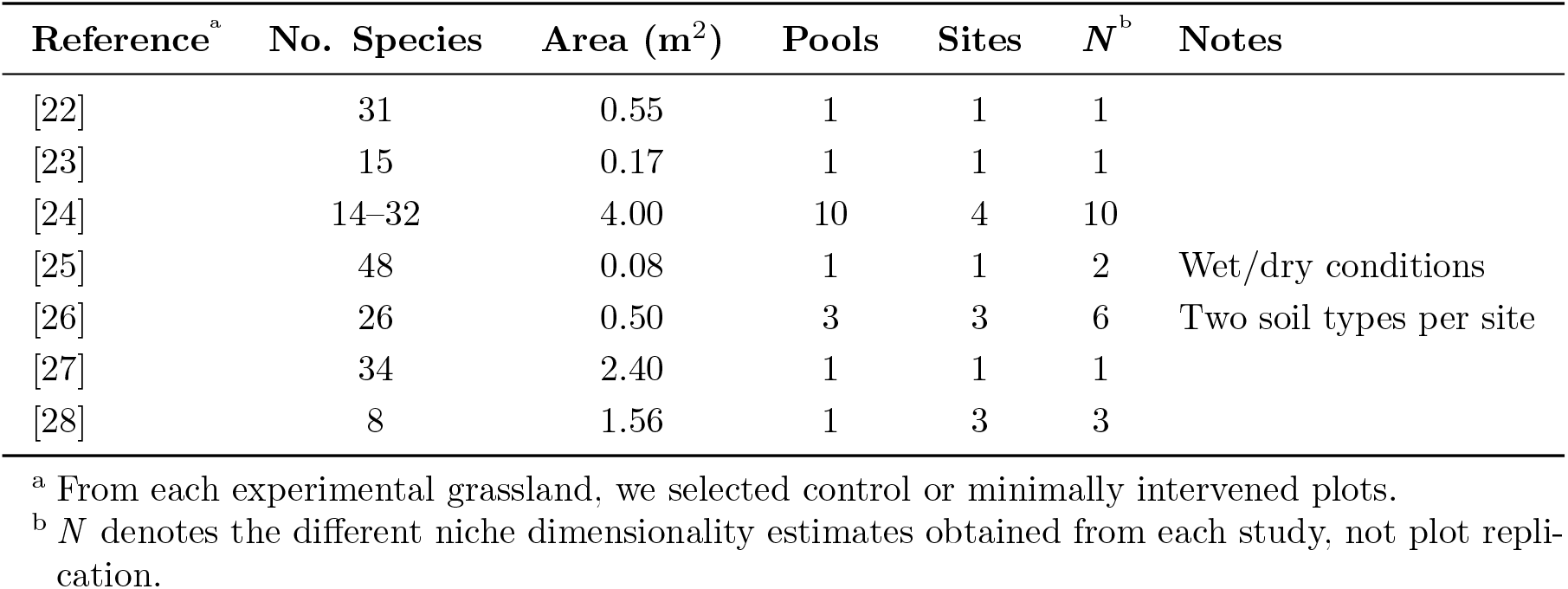
Experimental Data Summary.

As other latitudes were lacking from the experimental selection, we resorted to an open-access database, sPlotOpen [29], to increase the coverage of other biomes. sPlotOpen provides the opportunity to examine multiple realizations of community assembly of specific plant communities with a good phylogenetic coverage across a wide range of climatic and ecological contexts, thereby addressing the limitations of regional or habitat-specific experimental data. We selected grassland plots with at least 10 replicates of the same area, latitude, longitude, elevation, and slope. This resulted in 59 unique instances, assumed to represent communities shaped by similar assembly processes. We defined the species pool for each instance as the complete list of species present across its plots. For comparison, we also examined trait dimensionality in forest plots. Using the same procedure as with the grassland plots, we obtained a total of 189 unique instances of forest plots.

### 2.3 Phylogenetic and statistical analyses

We used R package U.PhyloMaker [30] under default settings to construct ultrametric phylogenies. We checked and corrected species names according to presence in the megatree. Those species absent from the megatree were included at the genus or family level. Organisms identified at the family level were also included in the phylogenies. In the case of the grassland plots, two of them included algae, so we excluded those from subsequent analyses. All further analyses were performed in R.

## 3 Results

First, we examined whether the richness of two datasets with measured species traits could be approximated with the measured species traits. We did so for [21] and [31], two datasets that included repeated measures of richness, full species lists, and species traits. We found that measured traits led to species richness lower than observed, stressing the need for considering more traits.

We estimated trait dimensionality in experimental grasslands, natural grasslands, and forests. We define trait dimensionality relative to the number of species in the pool as *γ*. Fig. 2 illustrates three examples of the values that dimensionality can adopt. Dimensionality adopted values that ranged from *γ <* 1, i.e., competition was driven by fewer traits than the number of species (lower panel, natural forests), to *γ* ≫ 1, i.e., competition was determined by many more traits than species (middle panel, natural grasslands), with intermediate cases where the number of traits determining competition were close to one per species, *γ* ≈ 1 (upper panel, experimental grasslands). The confidence intervals for dimensionality are tied to the variability on species richness for each example, and are relatively high.

**Figure 2.**
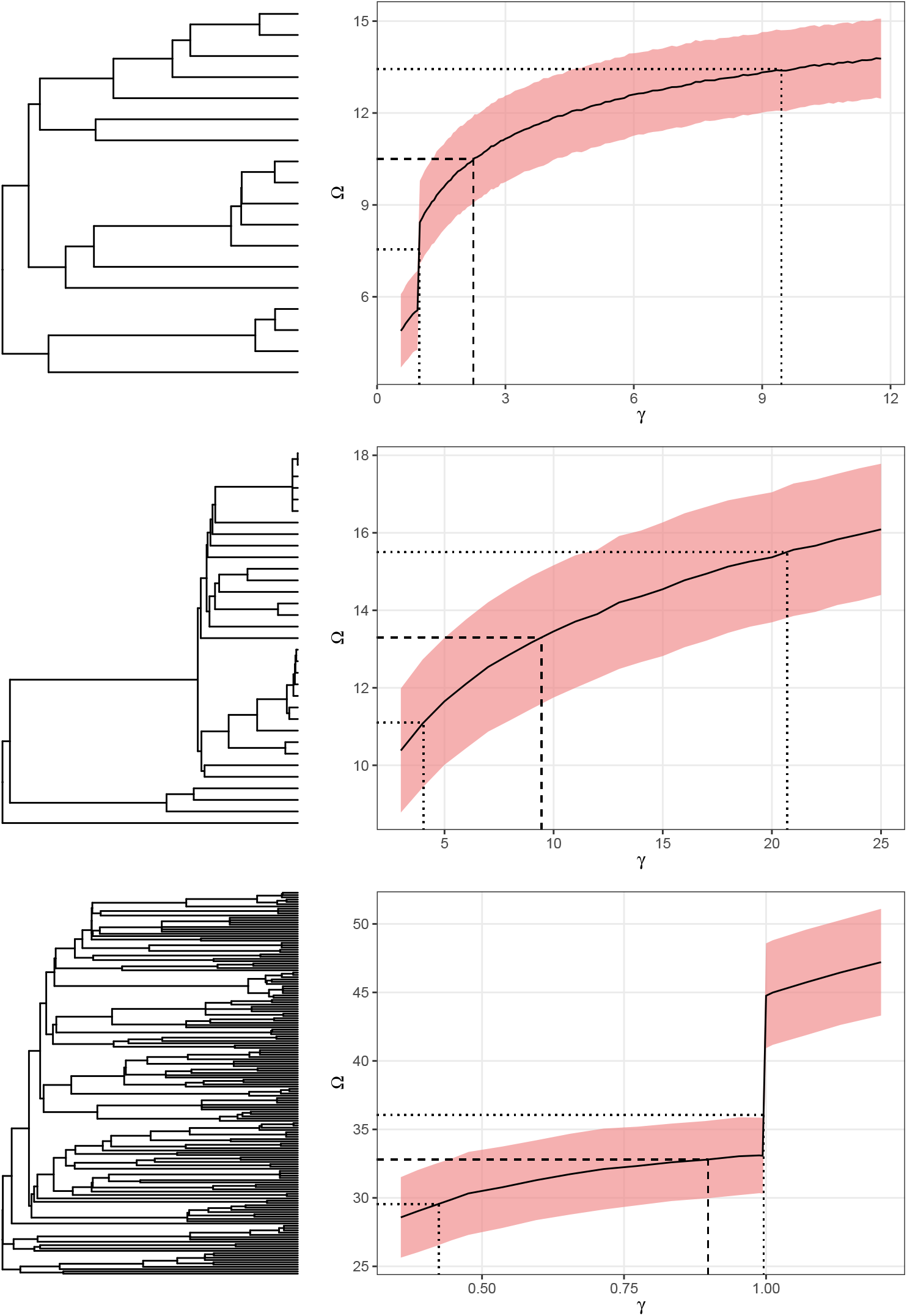
Trait dimensionality for three example communities. *Upper*, grassland community assembly experiment [24] using mix 107, which contains 18 species. The phylogenetic tree has a mix of early and late branching events, which produces a trait dimensionality above 2. *Middle*, natural grassland community, with 33 species. The phylogenetic tree denotes high shared evolutionary history, so trait dimensionality needs to be high to allow for the observed coexistence. *Bottom*, natural forest community with 168 species. Early branching events and low shared evolutionary history produce a trait dimensionality lower than 1, that is, less traits than species allow for the observed coexistence. Ribbons indicate ±1 s.d. around the mean estimate of richness for a given dimensionality.

The three types of communities examined in this work display a similar range of dimensionality values (Fig. 3). The mean value of dimensionality was not different between groups (Welch’s one way test, n.s.), although statistical power was very low (19-25%) and the distribution of forest dimensionality values seemed different to the grassland ones. In the case of experimental grasslands, 10% of the communities show a dimensionality below 1, and its mean value was 10.06. For the natural grasslands, 21.6% had trait dimensionality below 1, while its mean was 6.98. In the case of the forest communities, dimensionality was below 1 in 15.3% of the cases, and their mean dimensionality was 5.54.

**Figure 3.**
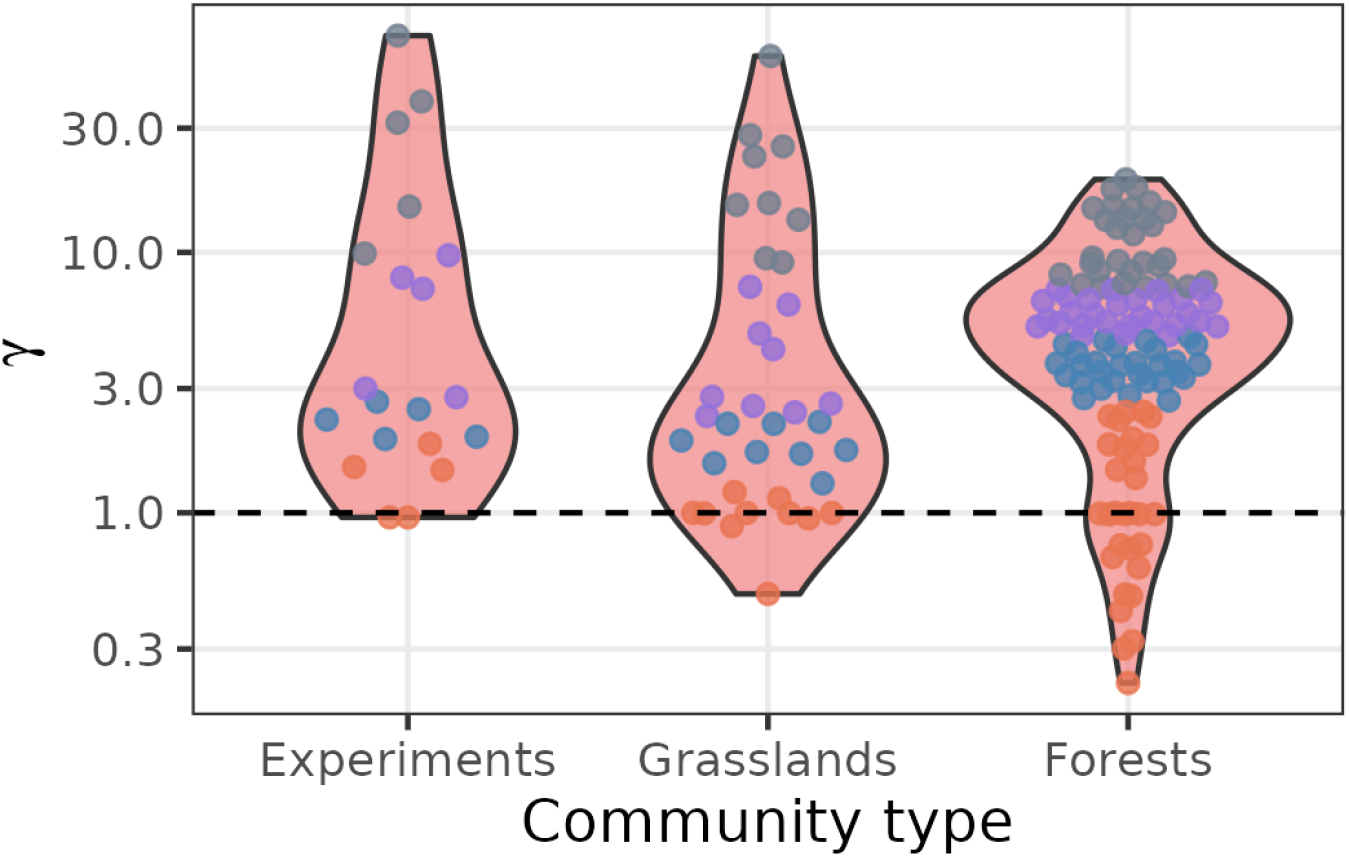
Trait dimensionality for experiments, natural grasslands, and forests. Equality of means can not be rejected at a .05 significance level, but statistical power was low.

Next, we looked at the effect of phylogenetic structure on dimensionality. We compared the dimensionality of communities with high and low phylogenetic similarity, expecting that for similar levels of survivorship dimensionality is higher in the case of phylogenetically close communities than for communities with distant communities. In general, communities with species more distant phylogenetically had a lower dimensionality (Fig. 4), evidenced with Wilcoxon rank sum tests (natural grasslands, *p*-value = 8.655 *×* 10^−7^; forests, *p*-value = 1.278 *×* 10^−10^). Experimental grasslands were not tested as *ρ* was low and homogeneous.

**Figure 4.**
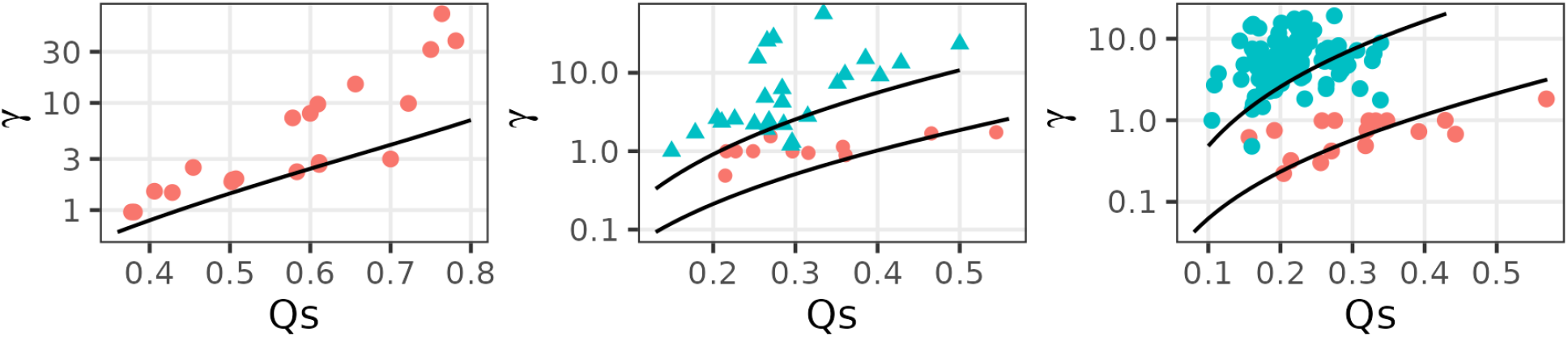
Dimensionality vs proportion of survivors, depending of mean *ρ* (color). Black line corresponds to the prediction of a theoretical community with phylogenetic distance (*ρ*) equal to the mean of the communities studied.

Finally, we examined whether a simplified phylogeny depicting mean field competition was able to predict accurately trait dimensionality estimated with full phylogenies. Indeed, *γ*^*\**^ and *γ* were highly correlated, almost in a linear fashion, for all three types of communities studied in this manuscript (Fig. 5).

**Figure 5.**
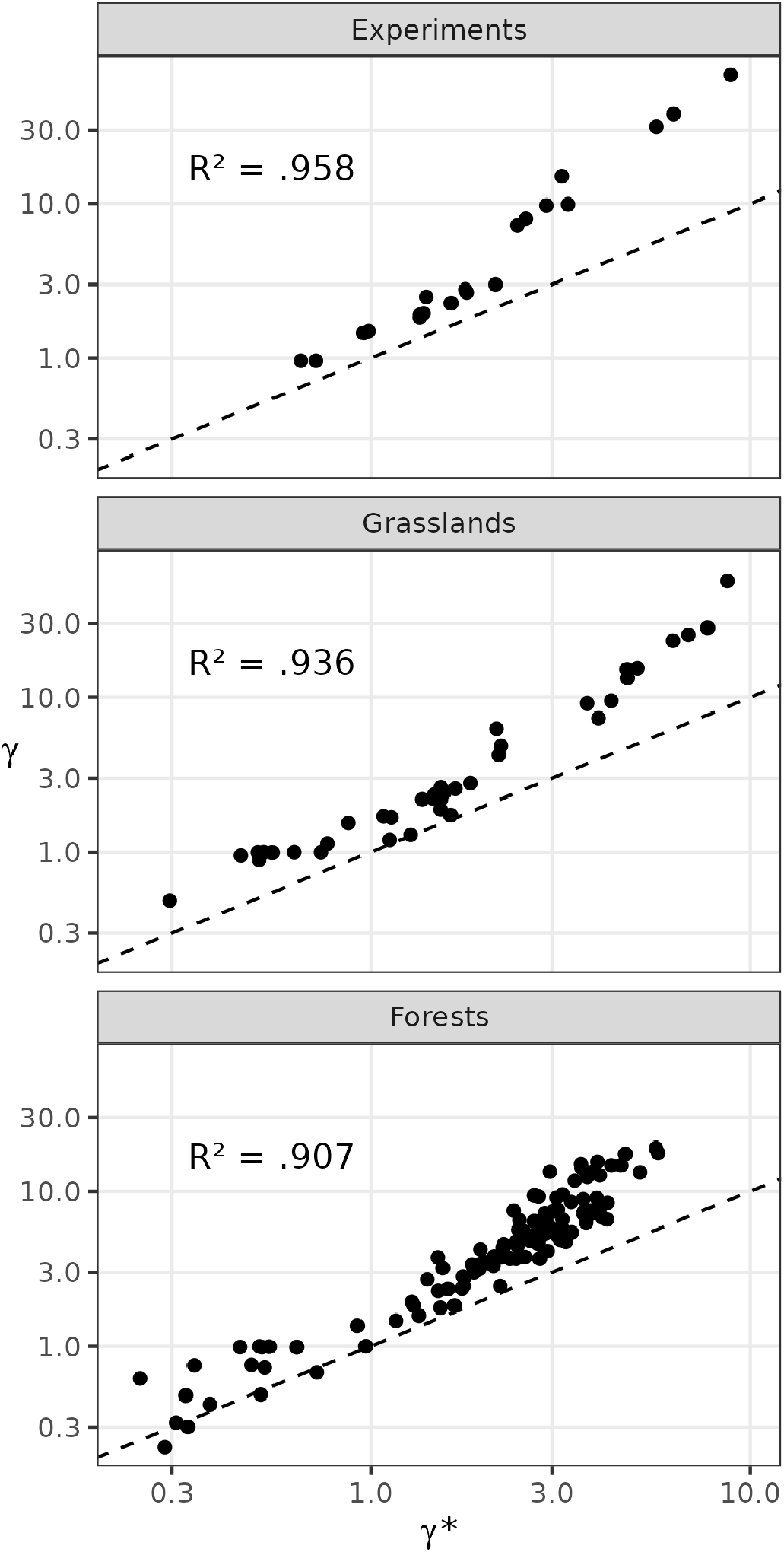
Dimensionality estimated from mean field competition predicts well the dimensionality obtained with full phylogenies.

As both types of grasslands show similar distributions, we hypothesized that they would respond in the same way to environmental drivers of dimensionality. They did so qualitatively (Tab. 2). The regression revealed complex interactions, yet the coefficient signs for different variables were consistent across experimental and natural grasslands. Put simply, increases in area tended to decrease dimensionality, while increases in the number of species tended to increase the traits per species needed to have a specific proportion of survivors.

**Table 2:** Regression Results for Grassland Dimensionality.

| Predictor | Experimental |  | Natural |  |
| --- | --- | --- | --- | --- |
| | Estimate ($\beta$) | S.E. | Estimate ($\beta$) | S.E. |
| (Intercept) | 136.80 | (58.50)* | 11.564 | (30.24) |
| Area | -192.26 | (31.03)*** | -1.398 | (1.15) |
| Species | -3.01 | (2.81) | -.503 | (.33) |
| Latitude | -2.72 | (1.16)* | -.384 | (.48) |
| Area × Species | 8.73 | (1.77)*** | .050 | (.02) |
| Area × Latitude | 4.20 | (.62)*** | .032 | (.02) |
| Species × Latitude | .06 | (.05) | .019 | (.01)** |
| Area × Species × Latitude | -.19 | (.04)*** | -.001 | (.0004)* |
| Adjusted $R^2$ | .87 | | .43 | |
| $p$-value | $1.31 \times 10^{-5}$ | | $6.09 \times 10^{-3}$ | |

Notes: Standard errors are in parentheses. Significance levels: \* $p < 0.05$ , \*\* $p < 0.01$ , \*\*\* $p < 0.001$ .

## 4 Discussion

One of the most striking findings of this study is that the number of traits required to explain species coexistence in both experimental and natural communities is higher than traditionally conceptualized. We found that trait dimensionality often exceeds one—that is, more traits are needed to explain coexistence under Lotka–Volterra dynamics than there are species. In two trait-measured datasets [21, 31], observed species richness exceeded that predicted by the available traits, implying that unmeasured traits are likely contributing to coexistence. Furthermore, we observed that communities composed of distantly related species, which likely have low niche overlap, tend to require fewer traits to explain coexistence across a range of occupancies compared to similar communities with closely related species. Despite this complexity, a simplified phylogeny—representing the mean-field competition case—explained a substantial portion of trait dimensionality and can serve as a lower bound for the number of traits needed to support observed community structure.

Current consensus is that a few number of traits (or combinations of those) is able to explain ecological strategies, life-histories, and, in turn, ecosystem functioning [11]. While the usefulness of this research program is undoubted at broad scales, several studies recognize the difficulty to apply it at local scales, where community assembly takes place [12, 32]. For example, trait combinations of plant communities are influenced at local scales by factors such as disturbance, soil conditions, niche partitioning and biotic interactions [6]. In fact, competition in ecological communities is structured by phylogeny, as shown recently with a similar model [16]. Here, we found that the number of traits that best explains richness in local communities, although varies wildly, is usually above the number of species in that community. This should not be surprising considering the competitive exclusion principle [33] and even Darwin’s own vision of competition.

We identified at least two limitations in our study. First, the lack of fitness differences in our model may represent a limitation of our work. Given the same phylogenetic tree, greater fitness differences lead to an effectively lower species richness. In that case, trait dimensionality would be underestimated by the procedure we developed in this study. Here, not only did we find a higher trait dimensionality than expected, but our method may also underestimate the true dimensionality of communities. Although the assumption of no fitness differences can be relaxed (see Appendix S6 in [17]), developing a framework to accurately estimate trait dimensionality when fitness differences are present is beyond the scope of the present study. A second limitation comes from the necessary requirements to estimate trait dimensionality. The estimation needs multiple replicates of assembly of the same species pool with a resolved phylogeny. The availability of community assembly experiments meeting these requirements is low and usually restricted to plant communities. This limitation may be overcome with observational studies, which face other caveats such as the determination of the species pool or the effect of heterogeneous conditions.

Here, we include a tentative examination of broad environmental factors that govern trait dimensionality in grassland communities. Our regression model for the experimental communities showed the effect of area, species, and latitude, and, interestingly, the signs are maintained for the natural communities, showing a similar but not as clear pattern as for the experimental ones. Conceptually, it is plausible that the area and number of species in which a community is packed affects trait dimensionality [34, 11, 35]. Latitude is also a long standing predictor of diversity patterns [36]. So, we demonstrated that by applying our approach to identify the effect of environmental conditions on trait dimensionality, we may expand the research programme of trait-environment studies to include the effect of phylogeny and competition. Also, the application of our approach to the design of experiments taking into account phylogeny may inform about the quantitative effects of multiple factors on coexistence.

The present research lays a foundation for further work. The approach that we have followed here can be modified to infer the phylogeny given the surviving species and the number of traits that determine their coexistence. In that way, we could compare infered and ‘true’ phylogenies to identify, *e*.*g*., wrongly resolved star-like radiations or incorrect molecular clock assumptions. Our framework might inform too about whether phylogeny is a good representative of species functions. Metagenomics of microbial communities has reached an state where it is possible to get well resolved genomes for an entire community, so instead of using the phylogenetic covariance matrix for the community to estimate coexistence or trait dimensionality, we could use a functional similarity matrix and assess which would fit better the observations. Actually, current research points to emergent microbial multispecies coexistence [37] with stable endpoints and functional convergence [38].

This study set out to understand trait dimensionality in ecological communities, approaching the subject from a theoretical standpoint that includes competition and niche differences. Understanding trait dimensionality as higher than previously acknowledged has profound implications, as many traits, and not only few, govern species coexistence. Future studies should explicitly explore the interaction of environmental variables and phylogeny on coexistence and trait dimensionality. Darwin’s intuition, the foundation of ecology and evolution, may still bear fruits.

## Acknowledgements

VJO was funded through grant PRIORITY (PID2021-127202NB-C22) awarded to JAC. Grant UNIQUE (PID2021-127202NB-C21) was awarded to D.A. Both grants were funded by MCIN/AEI/10.13039/501100011033 and “ERDF. A way of making Europe”.

